# Analysis of ancient human mitochondrial DNA from Verteba Cave, Ukraine: insights into the origins and expansions of the Late Neolithic-Chalcolithic Cututeni-Tripolye Culture

**DOI:** 10.1101/217109

**Authors:** Ken Wakabayashi, Ryan W. Schmidt, Takashi Gakuhari, Kae Koganebuchi, Motoyuki Ogawa, Jordan K. Karsten, Mykhailo Sokhatsky, Hiroki Oota

**Affiliations:** Department of Anatomy, Kitasato University, 1-15-1 Kitsasato, Sagamihara, Kanagawa 252-0374, Japan; School of Archaeology, Earth Institute, University College Dublin, Belfield, Dublin 4, Ireland; Kanazawa University, Center for Cultural Resource Studies, Kakuma, Kanazawa, Ishikawa 920-1164, Japan; Department of Anthropology and Religious Studies, University of Wisconsin-Oshkosh, Oshkosh, WI 54901; Borschiv Regional Museum, Ministry of Culture and Arts, Shevchenka St. 9, Bilche Zolote, Ukraine

**Keywords:** mtDNA, ancient DNA, Tajima’s D, Tripolye, Eneolithic, Ukraine

## Abstract

**Background:** The Eneolithic (~ 5,500 yrBP) site of Verteba Cave in Western Ukraine contains the largest collection of human skeletal remains associated with the archaeological Cucuteni-Tripolye Culture. Their subsistence economy is based largely on agro-pastoralism and had some of the largest and most dense settlement sites during the Middle Neolithic in all of Europe. To help understand the evolutionary history of the Tripolye people, we performed mtDNA analyses on ancient human remains excavated from several chambers within the cave.

**Results:** Burials at Verteba Cave are largely commingled and secondary in nature. A total of 68 individual bone specimens were analyzed. Most of these specimens were found in association with well-defined Tripolye artifacts. We determined 28 mtDNA D-Loop (368 bp) sequences and defined 8 sequence types, belonging to haplogroups H, HV, W, K, and T. These results do not suggest continuity with local pre-Eneolithic peoples, but rather complete population replacement. We constructed maximum parsimonious networks from the data and generated population genetic statistics. Nucleotide diversity (π) is low among all sequence types and our network analysis indicates highly similar mtDNA sequence types for samples in chamber G3. Using different sample sizes due to the uncertainly in number of individuals (11, 28, or 15), we found Tajima’s D statistic to vary. When all sequence types are included (11 or 28), we do not find a trend for demographic expansion (negative but not significantly different from zero); however, when only samples from Site 7 (peak occupation) are included, we find a significantly negative value, indicative of demographic expansion.

**Conclusions:** Our results suggest individuals buried at Verteba Cave had overall low mtDNA diversity, most likely due to increased conflict among sedentary farmers and nomadic pastoralists to the East and North. Early Farmers tend to show demographic expansion. We find different signatures of demographic expansion for the Tripolye people that may be caused by existing population structure or the spatiotemporal nature of ancient data. Regardless, peoples of the Tripolye Culture are more closely related to early European farmers and lack genetic continuity with Mesolithic hunter-gatherers or pre-Eneolithic groups in Ukraine.

## Background

Profound cultural transitions accompanied the Neolithic and Bronze Age in Europe. These transitions were catalyzed initially through large-scale migrations of agriculturists from the West Asia (modern day Anatolia) during the Neolithic, and later through the migrations of steppe herders from the Pontic-Caspian and Volga regions of modern Ukraine during the Bronze Age [1–4]. A continuing question of debate among researchers is how these migrations affected the genetic composition of modern-day Europe [5, 6]. A key question in this debate is whether these migrations that brought about cultural changes were the results of movements of people (demic diffusion model), as opposed to the movement of ideas and artifacts (cultural diffusion model) [7]. That is, were these large-scale migrations a process of cultural diffusion, with little genetic admixture among early Neolithic farmers and Mesolithic hunter-gatherers? Or, is a model of demic diffusion [7–10] more appropriate, whereby different regions of Europe were more or less affected by admixture with early farmers, and later steppe herders during the Bronze Age?

The transition to farming from a foraging lifestyle first appeared in the Near East around 10,500 years before present (yrBP) in modern-day southeastern Anatolia and Syria. Archaeologists have described two major and contemporaneous routes of expansion, namely the Continental (Danubian) and Mediterranean route. By 9,500 yrBP, farming spread into parts of Central Europe through the migration of peoples associated with the Linear Pottery culture (or *Linearbandkeramik*, LBK). These LBK cultures originated in Hungary and Slovakia (the Carpathian Basin) and then spread rapidly as far as the Paris Basin and Ukraine. A lingering question among archaeologists has always been whether these first farmers were descendants of local hunter-gatherers or whether they migrated from the Near East. Paleogenetic studies have generally suggested these early farmers were migrants, though in some places peoples continued to admix after the adoption of agriculture [5, 11–14.]

The region of Southeastern Europe (SE) has not been as extensively investigated as Southwestern (SW) Europe, Central Europe, or Southern Scandinavia, in terms of their ancient DNA variation. In contrast to Central Europe, the area of what is modern Ukraine saw the adoption of agriculture late. Although features of the Neolithic package are visible in Ukraine as early as 8,500–7,500 years before present (we use calBP to indicate radiocarbon dating), agriculture was not adopted as a primary subsistence economy until the Eneolithic or Chalcolithic period (around 5,500 yrBP) [15]. Whereas Central Europe saw mostly the process of demic diffusion [8], SE Europe seems to have adopted agriculture through innovative subsistence strategies through a transfer of ideas, little genetic influence, and genetic continuity from the Mesolithic to the Neolithic. Therefore, the Neolithic transition occurred at a slower pace, thus perhaps shaping the genetic composition of this region differently than in other parts of Europe.

Following the establishment of farming communities in the Balkan Peninsula, a series of complex societies formed, culminating in large settlements. By 5,500 yrBP, agriculture had reached Eastern Europe, in the form of the Cucuteni-Trypillian (C-T) complex in the area of present day Moldova, Romania, and Ukraine. This culture spanned close to 2,000 years and influenced much of SE Europe and the Baltic regions. It is known for elaborate anthropomorphic and animal figurines, as well as distinct elegantly painted pottery. Around 4,000 yrBP, these societies began to change, with the large settlements being abandoned, and archaeological evidence suggesting contact with nomadic steppe populations from the East.

We address the complex process of Neolithisation [7] in SE Europe by examining an Eneotlithic (Chalcolithic) Cucuteni-Trypillian site from Ukraine. For brevity, we will use the term Tripolye, as this is what it is known in Ukraine. Tripolye culture (7,100 – 5,000 calBP) is defined as Eneolithic based on the presence of copper artifacts and the onset of metallurgy, and ends at the beginning of the Bronze Age. The Tripolye culture occupied a large area from the Carpathian Mountains in the west to the Dnieper River in the east, and extended as far south as the Black Sea and north to Kiev (Fig. 1). Relative and absolute chronologies divide the Tripolye Culture into several phases (see Additional file 1), which normally accompany changes in pottery manufacture and decoration [16]. People associated with this culture are known as Trypillians.

**Figure 1.**
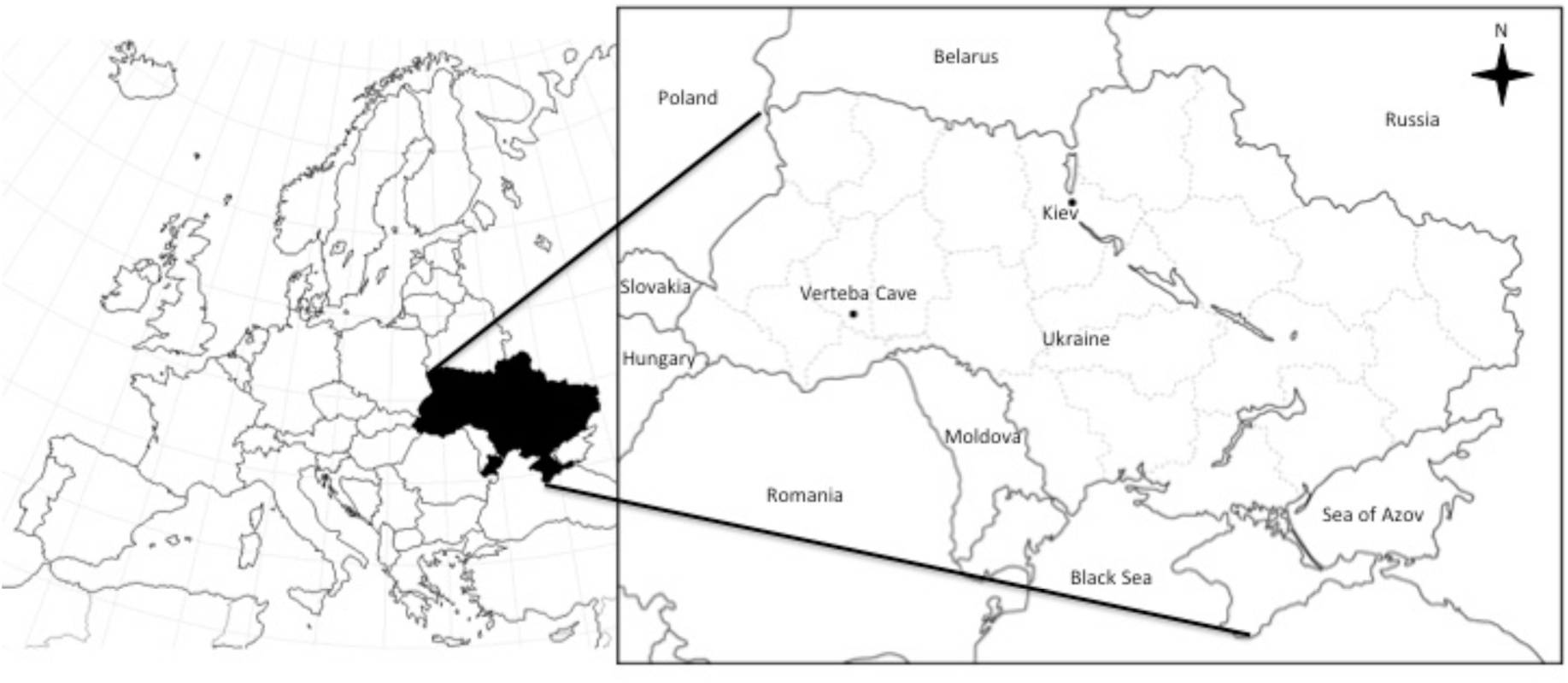
Map showing location of Verteba Cave.

In the present study, we investigate human remains found at a single Tripolye site known as Verteba Cave (VC). The occupation of the cave spanned over 1000 years, making it ideal to investigate the origins and possible genetic changes through time in the Trypillian people. Verteba Cave has been excavated as an archaeological site since the 1820s, though more intensive excavation has been ongoing since 1996 under the direction of co-author M. Sokhatsky (Borschiv Regional Museum of the Ukrainian Ministry of Culture and Arts). Verteba is a gypsum cave located in Western Ukraine, in the boreal forest-steppe zone of the East European Plain. It is one of many vast underground cave systems in the region.

Interpretations surrounding the use of the cave vary during these periods. Some believe the cave was used as a temporary shelter, while increasing archaeological evidence suggests use as a ritual site or a mortuary function. There is also evidence to support the idea that individuals buried in the cave, which are largely secondary in nature, are victims of warfare or sacrifice, due to the high frequency of blunt force trauma [17, 40]. Regardless, the cave contains the largest accumulation of human remains associated with the Tripolye culture. In fact, very few Tripolye culture human remains exist, making the cave one of the most important sites to investigate diet, health, pathology, and population history of Trypillian peoples [18, 19].

Using data obtained from the mitochondrial HVR-I region, we ask several interrelated questions about individuals buried at Verteba Cave. First, is there evidence for a maternal genetic continuity, and thus a degree of cultural diffusion, with local Mesolithic hunter-gatherers, as suggested in several recent studies? If there is not evidence for this, how are these individuals related to earlier farmer groups from SE or Central Europe? Is there any indication of a steppe influence for the Trypillian peoples? And lastly, can we infer something about Trypillian people’s expansion or collapse using maternal lineages? Several studies [20-22] have briefly addressed two of these three questions, indicating a link to early Neolithic farmers with some possibility for a link with Mesolithic hunter-gatherers. However, in those studies, the sample sizes tended to be smaller and the human remains used came from only a single chamber. Here, we analyze mtDNA data from several chambers in different locations found throughout the cave in an attempt to answer these questions.

## Results

To test reproducibility, we cut each specimen into 2 or 3 pieces and extracted DNA from each, and obtained 156 DNA extractions from 63 specimens (53 bones and 10 teeth) from chambers in the VC cave (Fig. 1). Using the DNA extractions as templates, we conducted PCR-amplification, and successfully amplified 38 specimens using all three defined overlapping primer-sets (Table 1). We sequenced 114 PCR amplicons using the Sanger method on an ABI 3130 Genetic Analyzer, and obtained sequences from 34 specimens for all three amplicons. We removed sequences from 6 specimens with amplicons that were mutually incompatible in nucleotide sequences, and finally obtained highly reliable 368-bp sequences from 28 specimens (Table 2, Additional file 2)

**Table 1.**
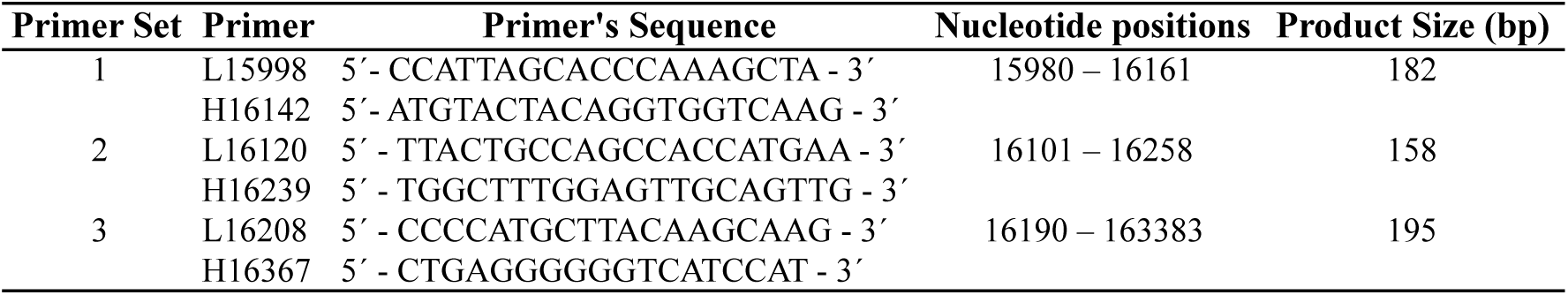
mtDNA Primers used in this study.

**Table 2.**
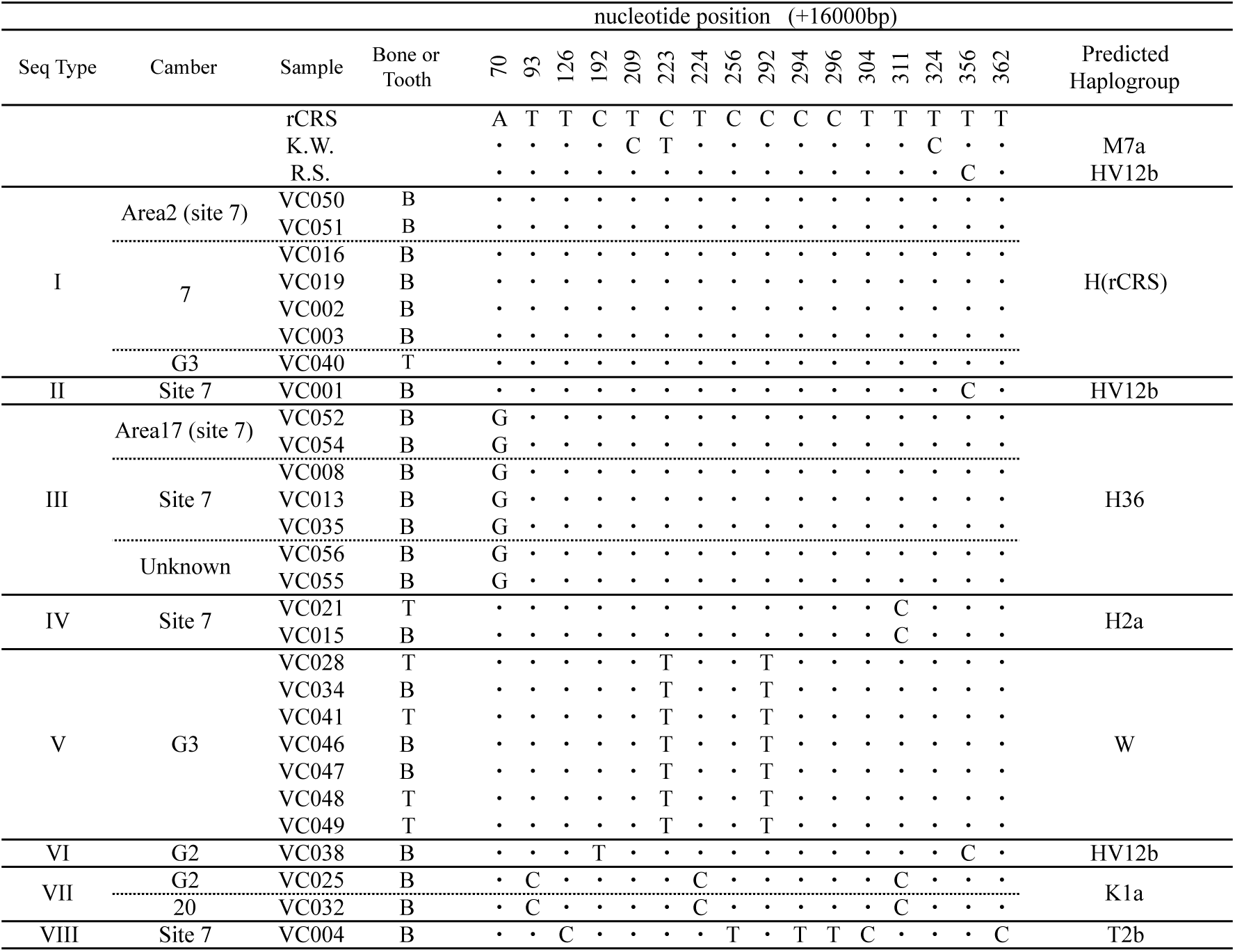
mtDNA nucleotide sequences of the Verteba Cave specimens

Eight defined sequence types (‘Seq Type’) of mtDNA D-loop were detected in 28 specimens, as well as the sequence types for two of us (K.W. and R.S.), who worked on DNA extraction. K.W. and R.S. sequence types were not identical to the eight sequence types except for Seq Type II from one bone specimen that was identical to the sequence type of R.S. (Table 2, Additional file 3). This sequence type corresponds to mitochondrial haplogroup HV12b, which is common in modern European populations. We could not eliminate the possibility that it is contamination, but included it in subsequent analyses. In addition to Seq type II, sequence type VI was predicted to be haplogroup HV. Sequence types I, III, and IV were predicted to be haplogroup H, while sequence types V, VII, and VIII were predicted to be haplogroups W, K, and T, respectively. Thus, all the haplotypes found in the VC specimens are common in modern European populations.

Based on the 8 sequence types of the mtDNA D-loop, a maximum parsimonious phylogenetic network was constructed (Fig. 2). Circles represent the sequence types, and the size of the circle is proportional to the number of samples. Numbers on the branches between the circles are nucleotide position numbers (+16,000) of the human mitochondrial genome sequence (rCRS [23]). Information about the location (chamber within the cave) where the specimen was excavated is also provided. Areas 2 and 17 are part of Site 7, and these are defined as a separate chamber, although they are located in close proximity within Site 7. The other chambers, Site 20, G2, and G3, are independent and separate locations within the cave. ‘Undefined’ chamber describes an unknown location within the cave.

**Figure 2.**
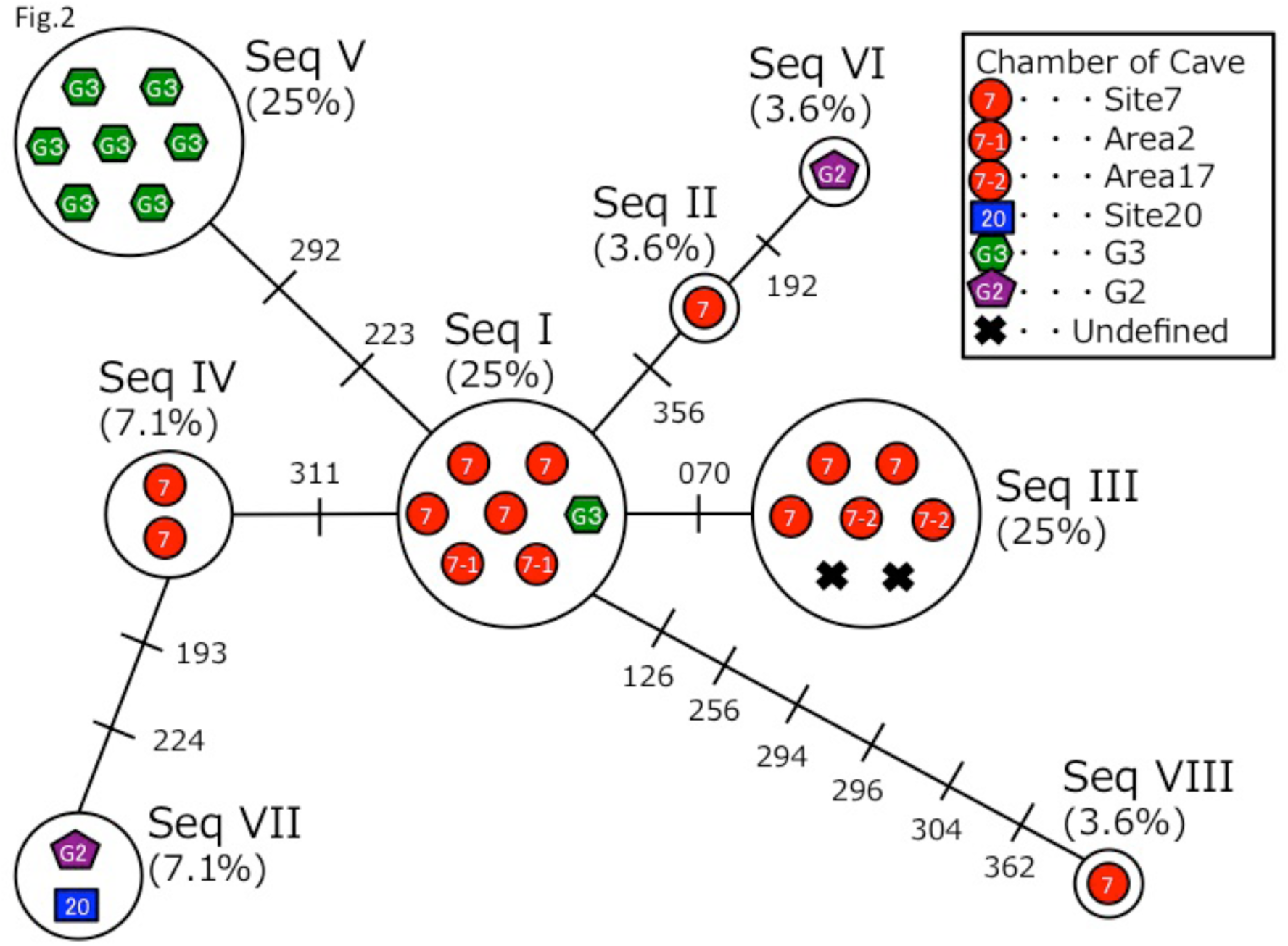
Maximum parsimonious network analysis. Circles represent the sequence types (‘Seq’), and the size of the circle is proportional to the number of samples. Numbers on the branches between the circles are nucleotide position numbers (+16,000) of the human mitochondrial genome sequence. Sequence types correlate to mtDNA haplogroups H, HV, W, K and T.

Specimens from each chamber showed deviation for the sequence type distribution observed in the sample set (Fig. 2). For example, specimens excavated from Site 7 had five unique sequence types, (I, II, III, IV, and VIII), while specimens excavated from chamber G3 had mainly one sequence type (V). For those specimens with the same sequence type, there are two possibilities: (1) the specimens belong to the same maternal lineage through several generations, or (2) the specimens (bone fragments and/or teeth) are from the same individual. This could be the case as all of the elements from site G3 were different (Size 28 in Tables 3, 4, 5). However, because archaeological excavation has not been completed for site G3, it is unclear which of these two possibilities is the case. We consider the two possibilities as both plausible. If the first possibility is correct, then we have 28 distinct individuals. If these lineages all come from the same individual, then we are left with only 11 individuals (Size 11 in Tables 3, 4, 5).

**Table 3.** Classification of geographical populations and genetic diversity statistics

| Area | Name of Population | Size | Number of Sequence Types | Number of Segregating Sites(S) | Nucleotide Diversity($\pi$) | Tajima's D |
| --- | --- | --- | --- | --- | --- | --- |
|  | **Verteba Cave** | **28** | **8** | **14** | **0.00634** | **-1.18985** |
|  |  | **11** | **8** | **14** | **0.00919** | **-1.28896** |
| West Europe<br>(N=595) | Basques | 45 | 27 | 31 | 0.00894 | -1.86317 * |
|  | British | 171 | 65 | 67 | 0.01189 | -1.95117 * |
|  | Danish | 31 | 25 | 28 | 0.01957 | -0.83873 |
|  | Germany | 107 | 73 | 59 | 0.01726 | -1.89550 * |
|  | Icelanders | 39 | 29 | 32 | 0.01404 | -1.30829 |
|  | Italy Toskany | 49 | 40 | 55 | 0.01343 | -2.05834 * |
|  | N_Spain | 30 | 26 | 35 | 0.01446 | -1.90059 * |
|  | Portugal | 54 | 37 | 39 | 0.01167 | -1.97766 * |
|  | Sardinian | 69 | 45 | 52 | 0.01180 | -2.05228 * |
| North Europe<br>(N=255) | Estonia | 26 | 23 | 32 | 0.01280 | -1.69668 * |
|  | Finns | 48 | 33 | 35 | 0.01045 | -1.83038 * |
|  | Norway | 82 | 17 | 20 | 0.00969 | -0.78572 |
|  | Sweden | 25 | 11 | 17 | 0.00909 | -0.98565 |
|  | Switzerland | 74 | 42 | 43 | 0.00971 | -1.94697 * |
| East Europe<br>(N=29) | Bulgaria | 29 | 22 | 39 | 0.01226 | -1.98753 * |
| West Asia<br>(N=71) | Anatolia Turks | 45 | 40 | 55 | 0.01482 | -2.03925 * |
|  | Turks | 26 | 24 | 52 | 0.01812 | -1.96694 * |
| Central Asia<br>(N=205) | Kazakh | 55 | 45 | 62 | 0.01764 | -1.87704 * |
|  | Kirghiz Sary Tash | 47 | 34 | 56 | 0.01613 | -1.95020 * |
|  | Kirghiz Talas | 48 | 43 | 55 | 0.01762 | -1.81413 * |
|  | Uighurs | 55 | 45 | 61 | 0.01602 | -2.04502 * |
| South Asia<br>(N=66) | Indian | 66 | 35 | 54 | 0.01614 | -1.59922 |
| Northeast Asia<br>(N=32) | Russia Siberia | 16 | 14 | 23 | 0.01478 | -1.10856 |
|  | Siberia Altai | 16 | 16 | 27 | 0.01513 | -1.47523 |
| East Asia<br>(N=593) | Cantonese | 18 | 18 | 36 | 0.01745 | -1.53414 |
|  | Changsha | 82 | 69 | 70 | 0.01821 | -1.92022 * |
|  | Mongolian | 103 | 82 | 79 | 0.01765 | -1.90508 * |
|  | Korean | 306 | 200 | 127 | 0.01458 | -2.32106 * |
|  | Xian | 84 | 76 | 63 | 0.01784 | -1.81700 * |

\* = $p < 0.05$

**Table 4.**
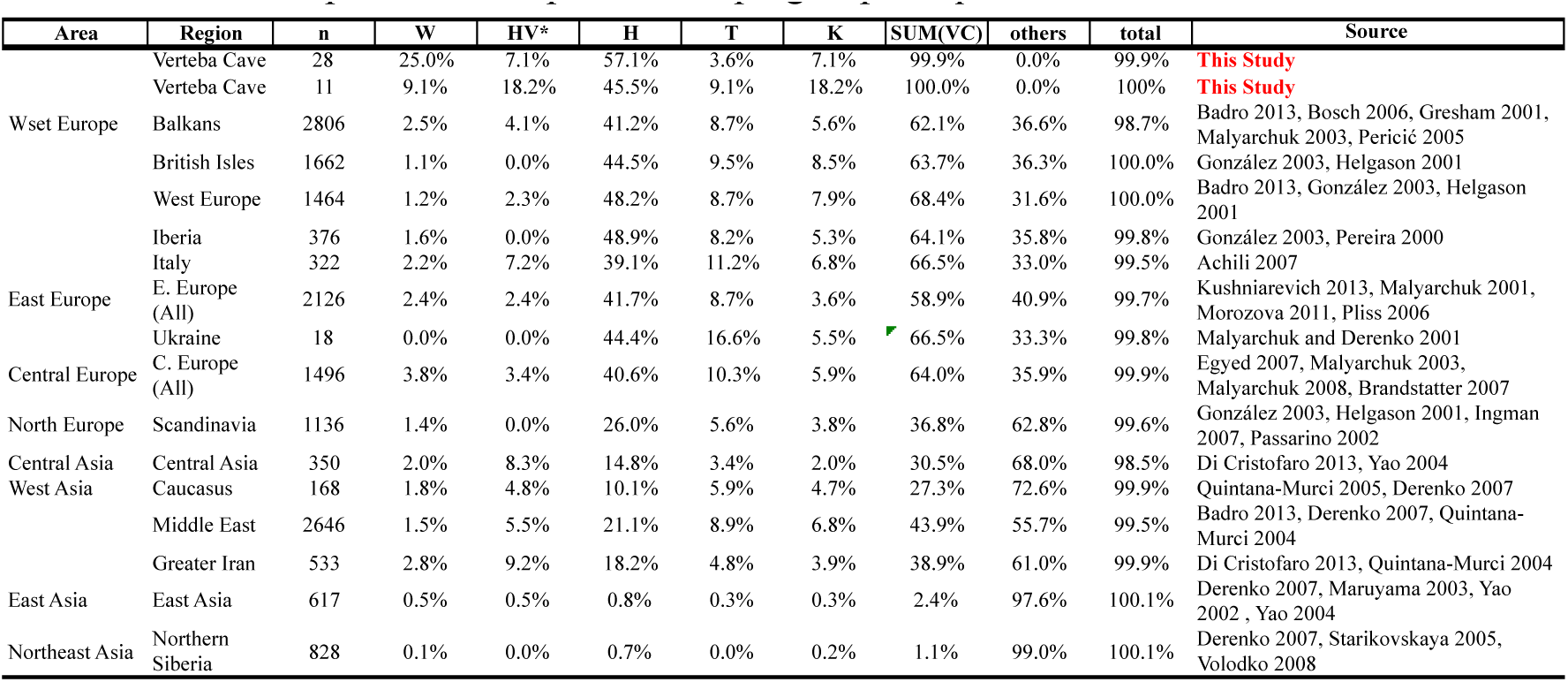
Modern Population Comparative Haplogroup Frequencies

**Table 5.**
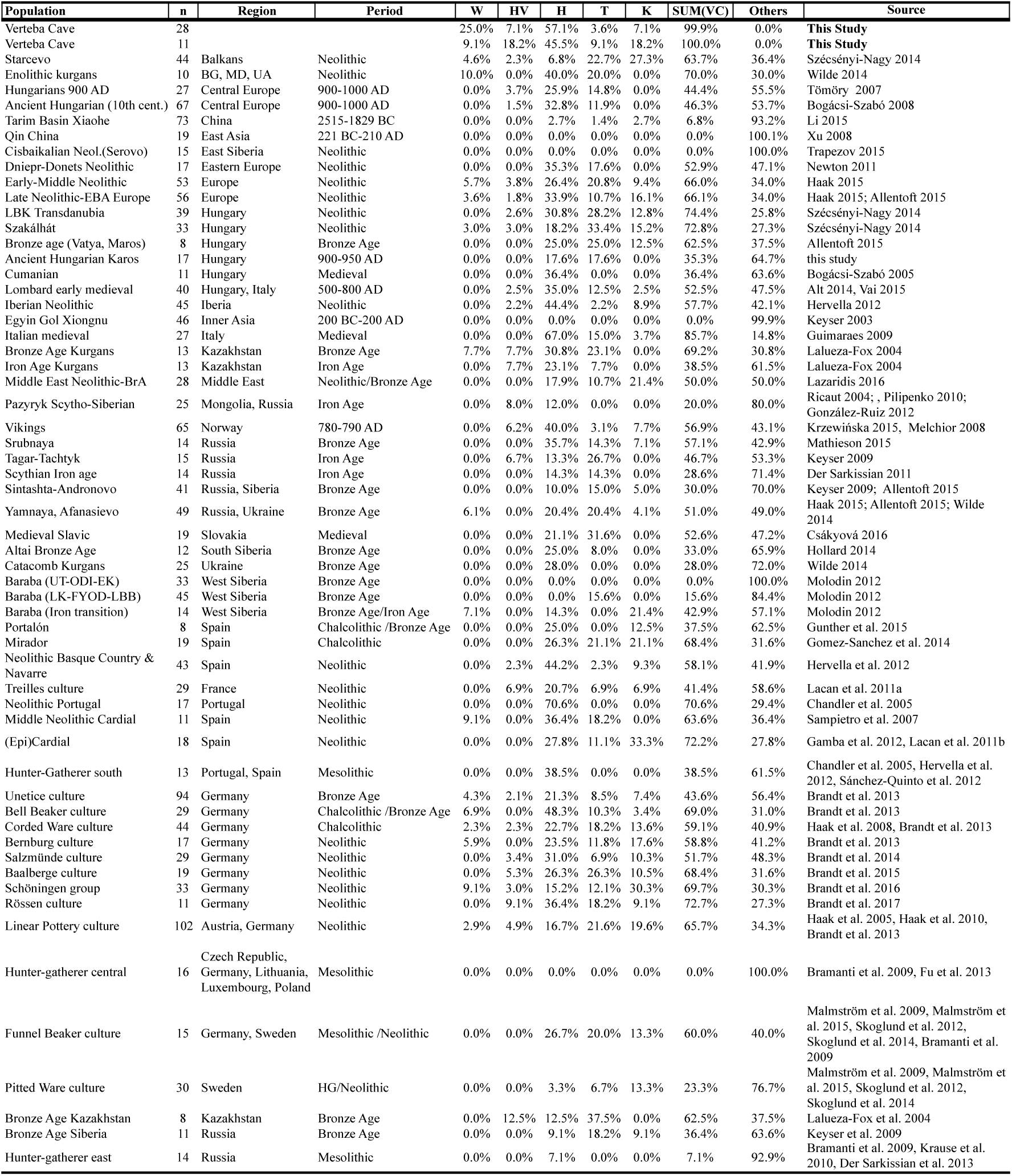
Ancient Population Comparative Haplogroup Frequencies

We next calculated population genetic statistics for the VC specimens (conducting analyses for both 28 and 11 individuals separately) and made comparisons with modern populations (Table 3). If we consider all the specimens belonging to a single population, then the number of sequence types, 8, was quite small compared to modern populations. Nucleotide diversity (π) in VC was 0.00634 when using 28 specimens as a population, and was 0.00919 when we only consider the total sample size to be 11. Both values of π are smaller than those of many modern populations, and relatively similar to those of Basques (0.00894), Norway (0.00969), Sweden (0.00909), and Switzerland (0.00971).

Tajima’s D statistic is a test of neutrality for nucleotide sequences, but when testing an obviously neutral locus (mtDNA), it suggests something about a population’s demographic history [24]. If Tajima’s D is statistically significant negative, then it suggests that the population has experienced demographic expansion. If Tajima’s D is not significantly different from zero, then it suggests that the population has maintained a constant population size [25]. Tajima’s D values of VC specimens were negative but not significantly different from zero. Thus, population genetic statistics of the VC specimens suggest that, if they are a contemporaneous population, then their genetic diversity was as small as modern Basques and Scandinavians who likely experienced bottleneck effects, and based on Tajima’s D test, the VC people were unlikely to be a population that experienced demographic expansion.

We compared the frequencies of the haplogroups predicted based on mtDNA D-loop sequences in the VC with those in the modern and the ancient populations from Europe and Asia (Table 4, 5). Haplogroup H was the most frequent (45.5 ~ 57.1%), and haplogroups W and HV* were the second most frequent (9.1 ~ 25.0% and 7.1 ~ 18.2%, respectively) in our samples. Haplogroup H is among the most common for modern Western, Central, and Eastern European populations, but the haplogroups W and HV* are relatively less frequent in modern European populations [26]. On the other hand, haplogroup H is less frequent in Central and West Asia, but slightly higher frequencies of haplogroup HV* are reported for those regions. In individuals from VC, haplogroups T and K also showed significant frequencies (3.6 ~ 9.1% and 7.1 ~ 18.2%, respectively). The ranges of the frequency of haplogroups reported here fall within the variation of modern European and West Asian populations (Table 4).

In comparison with ancient groups, VC haplogroup frequency is most similar to Neolithic farmer groups from Central Europe, as well as Funnel Beaker groups from Central (Salzmünde, Baalberge Cultures from Germany) and Northern Europe, particularly with respect to haplogroups H, HV, and T (Table 5). The Funnel Beaker cultural complex overlapped in territory with the Tripolye culture in the upper parts of the Dniester River and adjacent areas, and there is evidence for archaeological contact between the two cultures [21].

## Discussion

In this study we have attempted to answer a number of related questions surrounding Trypillian population history using data gleaned from maternal ancestry. We include at most 28 and a minimum of 11 individuals buried at Verteba Cave, an Eneolithic site associated with the Tripolye culture. Though we understand the limitation in using such data, we suspect our findings answer (in part), some fundamental aspects of Trypillian people’s genetic affiliation with other Neolithic groups.

To address the complexities associated with the transition to farming (or the process of Neolithisation), a wealth of ancient mtDNA sequences has been amassed from the Late Mesolithic to the Late Bronze Age [5, 27]. To explain these genetic changes in various regions of Europe, it is important to fully understand the genetic substratum spanning the Mesolithic-Neolithic transition. Studies have suggested that the maternal signature of local hunter-gatherer groups in many parts of Europe was homogenous with a relatively small population size and haplogroups dominated by lineage haplogroup U, such as U2, U4, U5a, U5b, and U8. Conversely, the genetic lineages of Early Neolithic farmers from Central and Southwest Europe are compromised by a wider array and diversity, with U lineages being less frequent. The mitochondrial Neolithic package arrived in Central Europe around 8,000 years BP with cultures associated with LBK farmers and the lineages are largely assigned to haplogroups N1a, T2, K, J, HV, V, W, and H. These lineages largely replaced local signatures of peoples associated with Mesolithic cultures, at least for areas of Central Europe [5].

We first explored whether individuals buried at Verteba Cave are more closely related to earlier Neolithic farmers from Central Europe, or perhaps have some connection with local hunter-gatherers, thus emphasizing the role of cultural diffusion in the adoption of agriculture. mtDNA haplogroup data for modern and ancient populations in Europe and West Asia have shown there was a discontinuity between late hunter-gatherers and early farmers, and later extant European populations, in most locations throughout Europe. An exception has been in SE Europe and the Baltic where cultural diffusion may have played a larger role [28].

The VC haplotype distribution indicates common haplotypes among Eurasian populations [26]. These include maternal haplogroups H, T, K, and W. The majority of haplotypes fall into haplogroup H, which is the most common haplogroup among modern-day Europeans and peoples of the West Asia, accounting for around 40% in Europeans [29] and is found in approximately 44% of modern Ukrainians [30]. This suggests that population continuity between the Tripolye and modern Ukrainians is a distinct possibility. Further, one individual displayed haplogroup T2b (Table 1). In addition to T2b being a possible marker of Anatolian expansion [5], it has also been found at high frequency in the Carpathian Mountains [31]. In one study [31], an individual from Bilche Zolote was found to have haplogroup T2b. Bilche Zolote is only 3 kilometers from the site of Verteba Cave, indicating some degree of local continuity with the Trypillian people.

Previous ancient DNA studies showed that hunter-gatherers before 6,500 yrBP in Europe commonly had haplogroups U, U4, U5, and H, whereas hunter-gatherers after 6,500 yrBP in Europe had less frequency of haplogroup H than before [5]. Haplogroups T and K appeared in hunter-gatherers only after 6,500 yrBP, indicating a degree of admixture in some places between farmers and hunter-gatherers. Farmers before and after 6,500 yrBP in Europe had haplogroups W, HV*, H, T, K, and these are also found in individuals buried at Verteba Cave (Table 5) [32]. Therefore, our data point to a common ancestry with early European farmers.

Our data also suggest population replacement. Mathieson et al. [42] analyzed a number of Neolithic Ukrainian samples (petrous bone) from several sites in southern, northern, and western Ukraine, dating to ~8,500 – 6,000 yrBP, and found exclusively U (U4 and U5) mtDNA lineages. It should be noted that ‘Neolithic’ in this context does not mean the adoption of agriculture, but rather simply coinciding with a change in material culture. They also analyzed several Trypillian individuals from Verteba Cave (different samples from the those included in this study). Similar to our findings, they found a wider diversity of mtDNA lineages, including H, HV, and T2b. These data, combined with our results, appear to confirm almost complete population replacement by individuals associated with the Tripolye Culture during the Middle to Late Neolithic.

Although we did not find any haplotypes common among hunter-gatherers (U lineage), there exists the possibility that some Trypillians inherited this haplotype from Mesolithic hunter-gatherers and thus some degree of cultural diffusion. In a study by Nikitin et al. [21] on mtDNA variation at Verteba Cave using teeth and cranial fragments, they found haplogroup composition similar to our results (mainly haplogroups H, HV, and T). However, Nikitin et al. [21] also found two individuals to have haplogroups U8 (U8b1a2 and U8b1b), which have been found among Paleolithic specimens [33]. Haplotype U8b1 has also been discovered in Neolithic Anatolian farmers [4]. Based on the data in [21], they were unable to distinguish whether their individuals with U8 were more similar to Paleolithic or West Asians from the Neolithic. Based on these data combined with our preliminary results, it appears the Trypillians were very much a distinct people who most likely displaced local hunter-gatherers with little admixture.

Haplogroup W was also observed in several specimens deriving from Site G3. Although we are unsure if all of these haplogroups come from a single or multiple individuals, this observation is interesting in that it is relatively rare and isolated among Neolithic samples. It has, however, been found in samples dating to the Bronze Age (Table 5) [35]. In the study by Wilde et al. [35], they found haplogroup W present in two samples from the Early Bronze Age associated with the Yamnaya and Usatovo cultures. The Usatovo culture (~ 3500 – 2500 BC) was found in Romania, Moldova, and southern Ukraine. It was the conglomeration of Tripolye and North Pontic steppe cultures. Therefore, this individual could link the Trypillian peoples to the Usatovo peoples and perhaps to the greater Yamnaya steppe migrations during the Bronze Age that lead to the Corded Ware Culture [36].

Verteba Cave contains archaeological evidence for the Tripolye cultural complex, including implements for agrarian cultivation, including grain processing. Based on the material culture, the immensity of certain settlements (some housing up to 10,000 people [34]), as well as the deterioration in biological health resulting from grain consumption, it is clear the subsistence economy of the Trypillians was reliant on agriculture. Population genetic statistics, in the form of Tajima’s D test, based on all VC mtDNA sequence data show that the Trypillian people (at least their maternal lineages) did not experience demographic expansion, at least during the Late Neolithic. Previous studies report that modern hunter-gatherers do not indicate a signal of demographic expansion in mismatch distribution and/or Tajima’s D test but farmers tend to show expansion based on increasing numbers of individuals living in sedentary conditions [37, 38].

Although we find a negative value for Tajima’s D when analyzing all of the sampled individuals (n=28), it is not significant. However, we are unable to treat our sample population as being contemporaneous. Although most individuals included in this study come from Site 7, which has been firmly established to date around 5,500 cal BP based on ceramics and radiocarbon dating [20, 39], other chambers in the cave are not as confidently dated. For example, chamber G3 had very few artifacts that came from the Tripolye culture. Given this caveat, we analyzed only individuals buried in Site 7 (n = 15) and found that Tajima’s D is negative (−1.50475) and significant (*p* = 0.04900). We theorize that this is due to the fact that individuals buried at Site 7 come from the time of peak occupation in the cave, and therefore were experiencing population aggregation and increased density, and therefore show a signal of demographic expansion. However, when we include individuals from other chambers (such as G3 that most likely date to a later period), we could be seeing the effects of population structure, which might cause the negative, but non-significantly different from zero of Tajima’s D indicating population size stability.

An alternative explanation for our observed population size stability when all individuals are included might be due to sampling strategy and the temporal component of ancient DNA sites. If a population migrates in low numbers into a new environment, such as the case with early migrating farmers from Anatolia, we would slowly see an increase in the population as they become increasingly sedentary over time. If we were to sample from this site and test for demographic expansion, we would most likely see that reflected in a statistic like Tajima’s D. As the population increases and resources reach an upper limit, groups would begin to fission and settle into new locations in close geographic proximity. Sampling individuals from each of these new sites, we would expect to see the maintenance of population stability since the groups, though genetically related, are spread out and thus would maintain population equilibrium with bi-directional migration. If, at some points in time, these sites again become aggregated because of increased population size, and we were to sample from this new, larger archaeological site, we would again witness demographic expansion simply because the population size has increased on the whole. Therefore, farmer populations as a whole (over the course of the Neolithic), generally see a trend for increased population expansion as local villages turn into larger settlements (as could be the case for peak occupation at Verteba Cave, where nearby settlements tended to be large). However, if we sample from each of those localities over time, as perhaps we are seeing with our results when we include samples from all sites (chambers), then we do not see demographic expansion, but rather maintenance of population size over time.

Archaeologically, it has been documented that Tripolye settlements began to disappear at the beginning of the Bronze Age. The reasons for this vary, but could be influenced by their interaction with steppe groups from the east. One of the possibilities for settlement abandonment is warfare. It has been well documented at Verteba that inter-personal violence was a common phenomena [17, 41]. Madden et al. [41] found a high degree of trauma-related cranial injuries among Trypillian burials. It is believed these individuals were killed by a raiding outside group and were later buried by members of the Tripolye culture. The Trypillians were one of the last of the “Old Europe” cultures that lived along the shores of the Danube River. By 5,800 calBP, many of these Neolithic Danubian cultures were wiped out after the arrival of pastoralists from the steppe [17]. It is believed that by 5,300 calBP, the Trypillians were in conflict with members of the Usatovo culture to the south, no longer benefitting from trade relationships across the forest-steppe boundaries. A reduction in mtDNA diversity, as shown by our results, could be explained through the targeted killing of both male and female Trypillians.

Another explanation for the sudden collapse of Tripolye culture, the crisis in Neolithic societies, and the observation of non-demographic expansion may be due to recent documentation of an early form of plague that was widespread from Siberia to the Baltic around 5,000 years BP [41]. Neolithic communities contracting this early form of the plague would have devastated Tripolye mega-sites, thus creating a demographic collapse that we are only glimpsing in our population genetic analyses. To get a better understanding of the demographic collapse in Tripolye society, we will need to obtain genome-wide data to further explore how these early agro-pastoralists eventually declined or were replaced by steppe nomads from the East.

## Materials and Methods

### Samples

Human remains for DNA analyses come from several excavation sites located within Verteba Cave (VC). VC is a mortuary site located outside the modern village of Bilche Zolote, Ternopil Oblast, Ukraine (Fig. 1). Most samples date to the Tripolye CII period (~5,500 BP) based exclusively on associated pottery found in the same cultural layer as the human remains (Additional file 1). Nikitin et al. [21] radiocarbon dated human and animal remains, as well as pottery sherds from Verteba and found the dates correspond to transitional phases in pottery decoration, with peak activity placed around 5,500 calBP. Almost all skeletal samples excavated within Verteba Cave are commingled and individual burials are difficult to identify. Therefore, in order to avoid sequencing the same individual twice, one of us (J.K.) collected second right metacarpal bones for analysis from a single chamber (Site 7). These samples were collected over several excavation field seasons (2008-2014). The remains were initially kept at the University of Wisconsin-Oshkosh, and later transferred to Kitasato University for processing. R.W.S. then collected bone and teeth samples on-site using sterile sampling methods (wearing coveralls, gloves, facemask) from four additional chambers (20, G1, G2, G3) during field seasons 2015–2016. These samples were directly deposited into sterile tubes and are now stored at University College, Dublin. A total of 63 individual human remains were analyzed for this study.

### Ancient DNA extraction

DNA extraction was carried out with ~100–300 mg of bone powder using a modified silica-column based protocol (Yang et al., 1998; Gamba et al., 2014; 2015). First, the surface of the bone was cleaned using a diamond drill bit a low speed. Then, we either used the drill bit to obtain powder, or powdered the bone in a mixer mill, ShakeMaster Auto ver.2.0 (BioMedical Science Inc.). For samples VC001-VC035 and VC046-VC063, we used the following protocol: bone powder was incubated for 24 h at 55°C followed by 24 h at 37°C in 2 ml tubes with 1 ml of lysis buffer in final concentrations of Tris HCL pH 7.4 20mM; Sarkosyl NL 0.7%; 0.5 M EDTA pH 8.0 47.5mM; Proteinase K 0.65U/ml, shaking at 300 rpm in a Thermomixer (Thermomixer comfort Eppendorf^®^). Samples were then centrifuged at 13,000 rpm for 10 m and the supernatant was removed. Fresh lysis buffer (1 ml) was then added to the pellet, vortexed, and the incubation and centrifugation steps were repeated.

The second supernatant was then transferred to an Amicon^®^ Ultra-4 Centrifugal Filter Unit 30K, diluted with 3 ml of TE and centrifuged at ~2,500 rpm until a final concentration of ~100 µl was obtained. This volume was then transferred to a silica column (MinElute PCR Purification Kit, QIAGEN) and purified according to manufacturers instructions, except at the final step adding TWEEN 20 (at 0.05% final concentration) to 60 µl EB buffer pre-heated to 60℃.

For samples VC036-VC044, our second protocol followed the first with the following modifications based on a “pre-digestion” step recommended in Gamba et al. (2015): Fresh lysis buffer was added to the powder and incubated in a Thermomixer for 1 h at 56°C shaking at 1200 rpm. After 1 h, the sample was centrifuged for 2 m at 13,000 rpm and the supernatant was discarded. Fresh lysis buffer was then added to the pellet and incubated at 56°C for 1 h followed by 37°C overnight, shaking at 1200 rpm. After ultra-filtration in an Amicon^®^ Ultra-4 Centrifugal Filter Unit 30K, the sample was transferred to a silica column (MinElute PCR Purification Kit, QIAGEN) with the following modifications: After adding the PB buffer, the centrifugation speed was reduced to 8,000 rpm and the elution step included a 5 m incubation at room temperature.

### Contamination Controls

To control for contamination, all pre-PCR procedures were conducted in a controlled-access, positive pressure laboratory with HEPA-filtered air that is installed exclusively for ancient DNA analysis at Kitasato University. Disposable protective clothing was worn during all sampling and extraction procedures. Pipettes with aerosol-resistant tips were used. The lab is cleaned prior to all procedures using DNA-OFF™ (Takara, Japan), and exposed to UV irradiation for at least two hours after each procedure. All PCR reactions and post-PCR procedures were performed in a separate laboratory. The movement of laboratory materials and personnel was always unidirectional, from the ancient to modern facilities. The mtDNA of all researchers with access to the clean lab were typed and compared with the results. At least two extractions and two amplifications were performed at separate times to assess the authenticity of our results.

### PCR amplification and direct sequencing

The hypervariable region I (HVRI) of the mitochondrial D-loop (367 base pairs, nucleotide positions 15999 – 16366) was targeted for amplification using three sets of overlapping primers (Table 1). PCR amplification was carried out using 2 µl extract in a 50 µl reaction mixture containing Ex Taq Hot Start, 10x Ex Taq buffer (containing MgCl_2_), 2.5 mM dNTPs, and 10uM each primer. PCR conditions were 94°C for 5 min followed by 40 cycles of 94°C for 30 sec, 55°C and 61.5°C for Primer Set 1and Primer Sets 2/3, respectively, annealing temperature for 30 sec, 72°C for 30 sec, extension of 72°C for 5 min, and a hold at 4°C. A negative control was included in each PCR run. PCR products were visualized on 2% agarose gel containing ethidium bromide, and purified using the MinElute PCR Purification Kit, QIAGEN. Amplification products were sequenced using Applied Biosystems Big Dye protocols and analyzed on an Applied Biosystems 3130 Genetic Analyzer at Kitasato University.

### Data analysis

Sequences were visually inspected, corrected and compared against the revised Cambridge Reference Sequence (rCRS) (Andrews et al. 1999) using DNASTAR^®^ Lasergene software (SeqMan Pro). Maximum-parsimonious phylogenetic network was constructed by hand. HVS-I haplotypes were classified into possible haplogroups and sub-haplogroups using MITOMAP (Lott et al. 2013). We compared the frequencies of haplogroups found in VC with ancient and modern people (58 populations / 14,735 individuals). Using the nucleotide sequence information obtained this time and the known nucleotide sequence information analyzed in [39] (29 populations / 1846 individuals), population genetic statistics, the number of sequence types, the number of polymorphic sites, the nucleotide diversity (π), Tajima's D, were calculated by software, DnaSP 5.

